# Extensive substrate recognition by the streptococcal antibody-degrading enzymes IdeS and EndoS

**DOI:** 10.1101/2022.06.19.496714

**Authors:** Abigail S. L. Sudol, John Butler, Dylan P. Ivory, Ivo Tews, Max Crispin

**Author notes:** Correspondence to Max Crispin and Ivo Tews.

## Abstract

Enzymatic cleavage of IgG antibodies is a common strategy used by pathogenic bacteria to ablate immune effector function. The *Streptococcus pyogenes* bacterium secretes the protease IdeS and the glycosidase EndoS, which specifically catalyse cleavage and deglycosylation of human IgG, respectively. IdeS has received clinical approval for kidney transplantation in hypersensitised individuals, while EndoS has found application in engineering antibody glycosylation. Here, we present crystal structures of both enzymes in complex with their IgG1 Fc substrate, which was achieved using Fc engineering to disfavour preferential Fc crystallisation. The IdeS protease displays extensive Fc recognition and encases the antibody hinge. Conversely, the glycan hydrolase domain in EndoS traps the Fc glycan in a flipped-out conformation, while additional recognition of the Fc peptide is driven by the so-called carbohydrate binding module. Understanding the molecular basis of antibody recognition by bacterial enzymes will facilitate the development of next-generation enzymes for clinical and biotechnological use.

## Introduction

The bacterium *Streptococcus pyogenes* has evolved a diverse range of mechanisms for evading the human adaptive immune system^1^. Infection with *S. pyogenes* can be mild, causing for example throat infections, but at the other extreme can cause terminal necrotising fasciitis^2^. Two enzymes secreted by this bacterium, IdeS^3^ and EndoS^4^, directly target and cleave IgG antibodies, and thereby impede cellular responses through immune recruitment mediated by the antibody Fc domain. The specificity of these enzymes for IgG has led to the development of a wide range of clinical and biotechnology applications^5^ and has warranted extensive studies of their enzymology.

Of the two immune evasion factors, IdeS is most advanced in clinical development. *S. pyogenes* expresses two variants of this enzyme (often distinguished by naming the first and second variants IdeS/Mac-1 and Mac-2, respectively), which display less than 50 % sequence identity within the middle third of the protein^6^, but nonetheless exhibit largely indistinguishable endopeptidase activity^7^. The enzyme targets IgG by cleaving within the lower hinge region, yielding F(ab′)_2_ and Fc fragments^3,8,9^, an activity which has enabled its development as a pre-treatment for transplantation in hypersensitized individuals with chronic kidney disease (Imlifidase, brand name Idefirix®)^10-12^. Along with EndoS, it has further potential use in the deactivation of pathogenic antibodies in autoimmune disorders^13-19^, deactivation of neutralising antibodies for *in vivo* gene therapy^20^, and for the potentiation of therapeutic antibodies by deactivation of competing serum IgG^21,22^. Imlifidase has also been used in combination with EndoS for inactivation of donor-specific antibodies in murine allogeneic bone marrow transplantation^23^.

The endoglycosidase EndoS has additional biotechnological applications in engineering antibody glycosylation^24^: it hydrolyses the β-1,4 linkage between the first two N-acetylglucosamine (GlcNAc) residues within biantennary complex-type N-linked glycans on IgG Fc, thereby removing the majority of the glycan^4^. The related enzyme EndoS2 from serotype M49 of *S. pyogenes* also targets IgG^25^ but exhibits broader glycan specificity^26^. Variants of both enzymes have been utilised in transglycosylation of various glycoforms to intact IgG to enable precise antibody glycan remodelling^27-29^.

It is still unclear, however, how exactly these enzymes specifically target and degrade IgG. Full cleavage of an antibody by IdeS occurs in two distinct steps, in which the second chain is cleaved more slowly^8,9^; this observation, along with the finding that IdeS exhibits low activity towards synthetic hinge peptides^30^, suggests a more extensive recognition interface with the target IgG. Similarly, multiple domains within EndoS contribute to substrate recognition and catalysis^31-33^, but the collective mechanism of these has not been fully resolved.

Here, we sought to understand the molecular basis behind the unique substrate specificity of these enzymes using X-ray crystallography. We identified and replaced the IgG Fc residue E382, which consistently forms salt bridge interactions in Fc crystal structures. This strategy discouraged Fc self-crystallisation and promoted crystallisation of the protein complexes. We present crystal structures of IdeS/IgG1 and EndoS/IgG1 complexes, to a resolution of 2.34 Å and 3.2 Å, respectively, and map the extensive interfaces that are formed in these complexes. Understanding substrate interaction and recognition will facilitate development as highly specific biotechnological tools and further clinical development.

## Results and Discussion

### Analysis of Fc crystal structures for Fc engineering

The co-crystallisation of IgG Fc with enzymes is notoriously difficult, due to the inherent ability of the Fc fragment to crystallise on its own. We therefore sought to identify favourable contacts present in typical Fc crystals, in order to devise a strategy for the elimination of its selective self-crystallisation.

We have observed, from looking at structures currently present in the PDB, that IgG Fc commonly crystallises in the *P*2_1_2_1_2_1_ space group (52 % of 125 structures, as of August 2022). We studied the crystal lattice contacts present in a typical, wild-type Fc structure (PDB code 3AVE), in order to identify amino acid residues which are important in this favourable packing arrangement. As calculated in PISA^34^, model 3AVE forms thirteen salt bridges and fifteen hydrogen bonds with neighbouring molecules within its crystal lattice (Fig. 1a). In addition, contacts are largely conserved across both Fc chains, resulting in a tight packing arrangement (Fig. 1a/b). We identified residue E382, which forms salt bridges with R255 in a neighbouring Fc molecule (and vice versa), in both Fc chains (Fig. 1b). We hypothesised that replacement of this residue would hinder the self-association of the Fc into this preferred crystal lattice, and therefore designed three IgG1 Fc variants: E382R, E382S and E382A, which we collectively term as “Fx” variants.

**Fig. 1:**
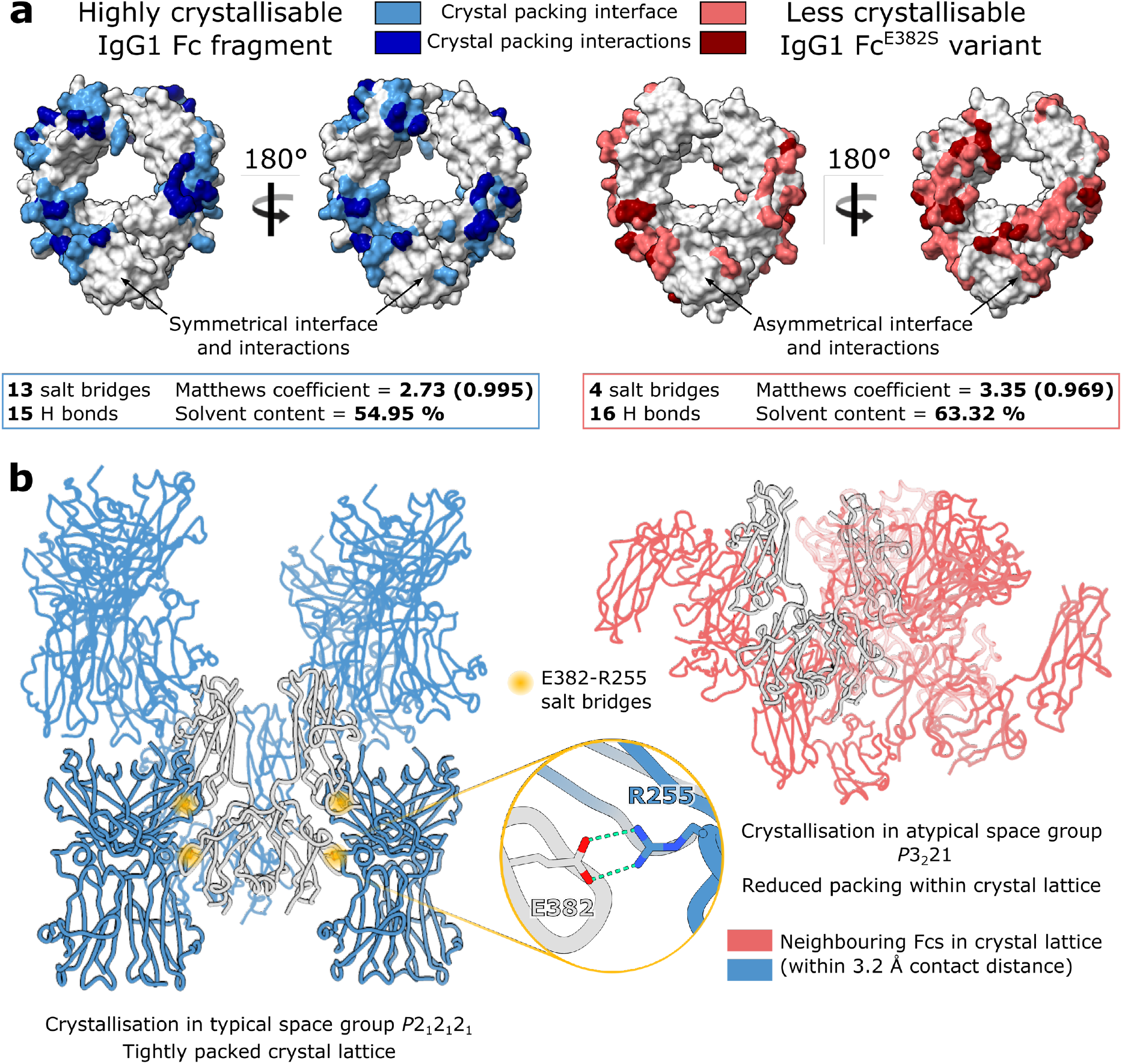
Observed crystal packing in wild-type and “less crystallisable” IgG1 Fc fragments. **a** Analysis of crystal packing interface and interactions present in a typical, wild-type IgG1 Fc crystal structure (PDB code 3AVE) and IgG1 Fc^E382S^ variant, as calculated by PISA^34^. **b** Crystal packing resulting from crystallisation in typical space group *P*2_1_2_1_2_1_ and atypical space group *P*3_2_21, for the wild-type IgG1 Fc and Fc^E382S^ variant, respectively. E382-R255 salt bridges between symmetry-related Fcs in the crystal lattice are highlighted with a yellow circle. Neighbouring Fcs in the crystal lattice contacting the origin Fc within a 3.2 Å contact distance are shown. Analysis relating to the wild-type Fc and Fc^E382S^ variant is depicted in blue and red, respectively.

We obtained crystals of the Fc E382S variant, which were found to have grown in an atypical space group *P*3_2_21. The structure was determined by molecular replacement using 3AVE as a search model and refined to a resolution of 3.04 Å (Supplementary Table 1). Analysis of the crystal contacts revealed that this variant makes fewer interactions with symmetry-related molecules in the crystal (four salt bridges and sixteen hydrogen bonds; Fig. 1a), which are asymmetrical across the two Fc chains, resulting in altered crystal packing (Fig. 1b). Furthermore, as calculated within the ccp4i2 interface^35^, the E382S variant had a higher solvent content and Matthews coefficient compared to the wild-type Fc (Fig. 1a), indicating that the molecules are less tightly packed in the variant crystal form. We conclude that these Fx variants are uniquely suited for screening attempts, as crystallisation of Fc fragments has been rendered less favourable; we subsequently used these variants for screening of enzyme-Fc complexes.

### The IdeS-IgG1 Fc complex

Using our panel of IgG1 Fx mutants, IdeS from *Streptococcus pyogenes* (strain MGAS15252), containing a C94A mutation to abolish catalytic activity, was crystallised in complex with IgG1 Fc (E382A variant), in space group *C*121 (Supplementary Table 2). The structure was determined by molecular replacement with 1Y08 and 3AVE search models. Electron density resolves amino acids 43-339 in IdeS, as well as 228-445 and 229-444 for chains A and B in IgG1 Fc, respectively. We additionally observe density for seven/eight monosaccharide residues at the N-linked glycosylation site (at N297) on Fc chains A and B, comprising a fucosylated biantennary glycan with a single β-1,2-linked GlcNAc on the mannose 6-arm (chain A) and the equivalent glycan with terminal β-1,2-linked GlcNAc on both arms (chain B). The final structure was refined to 2.34 Å (Supplementary Table 2) and is depicted in Fig. 2.

**Fig. 2:**
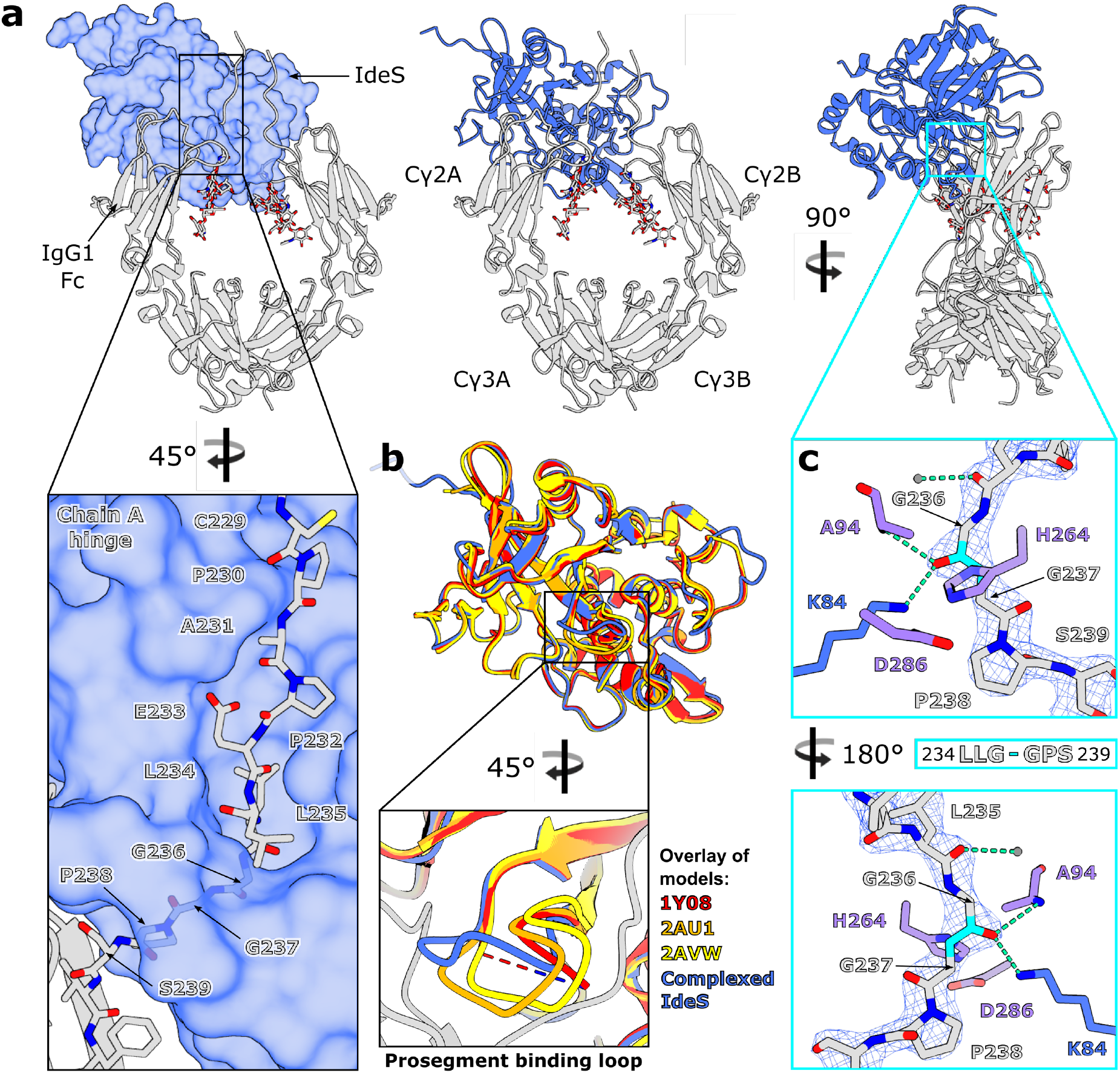
Crystal structure of IgG1 Fc^E382A^-IdeS^C94A^ complex. **a** Overall structure of complex, with IdeS shown as a surface and IgG1 Fc shown as a cartoon. N-linked glycans and the Fc hinge peptide in the focussed panel are shown as sticks and coloured with oxygen, nitrogen and sulphur atoms in red, blue and yellow, respectively. **b** Superposition of complexed IdeS with three published apo structures of IdeS (PDB codes 1Y08, 2AU1 and 2AVW, coloured in red, orange and yellow, respectively) and focused view of the prosegment binding loop. **c** Binding of Fc hinge peptide within the IdeS active site. Fc peptide and IdeS active site residues are depicted as sticks and coloured by heteroatom; catalytic triad residues are coloured purple. The scissile peptide bond is coloured in cyan. Hydrogen bonds are depicted as green dashes; catalytic water is depicted as a grey circle. The final 2F_obs_-F_calc_ electron density map corresponding to the Fc peptide is shown (weighted at 1.5 σ). **a**,**b**,**c** IdeS is coloured blue; IgG1 Fc is coloured in silver.

#### Overall structure of IdeS-IgG1 Fc complex

The crystal structure shows asymmetric binding of IdeS across the Cγ2 domains of the Fc (Fig. 2a). Although the crystalline state of a molecule doesn’t necessarily represent its biologically-relevant form, the combination of evidence from crystallography (Fig. 2a), analytical size exclusion chromatography (Supplementary Fig. 1) and previous kinetic analyses^36^ allows us to conclude that IdeS functions predominantly in a monomeric form. The enzyme appears to clamp down over the lower hinge region of one Fc chain (Fig. 2a), creating a cavity in which the catalytic residues are brought into close proximity with the cleavage site. Consequently, the Cγ2 domain in chain A is pulled away from chain B; this is reflected in a greater root mean squared deviation between Cαs in the Cγ2 domains (1.347 Å compared to 0.675 Å in wild-type Fc 3AVE, calculated in ChimeraX^37^ for residues 237-341) and higher atomic B factors in this domain (Supplementary Fig. 2a).

#### Role of prosegment binding loop in complex formation

IdeS crystallised in complex with IgG1 Fc here is the Mac-2 variant, and thus displays sequence diversity against the three published apo structures of IdeS (all of which are the Mac-1 variant; Supplementary Fig. 3a). Despite this, a structural alignment shows very few deviations (Fig. 2b). Complexed IdeS contains ten alpha helices and twelve beta strands, as calculated by DSSP^38,39^ (Supplementary Fig. 2b); we note that the loop located between beta strands seven and eight is modelled in distinct conformations for each of the apo structures and is not included within 1Y08^40^ (Fig. 2b), signifying its inherent flexibility in the apo form. This loop is equivalent to the “prosegment binding loop” present in other papain superfamily cysteine proteases; in these enzymes, which are synthesised as inactive zymogens, this loop packs against the prosegment as a mode of inhibition^41-43^. In complexed IdeS, the loop curls upwards to accommodate the Fc hinge within the active site cavity (Fig. 2a).

**Fig. 3:**
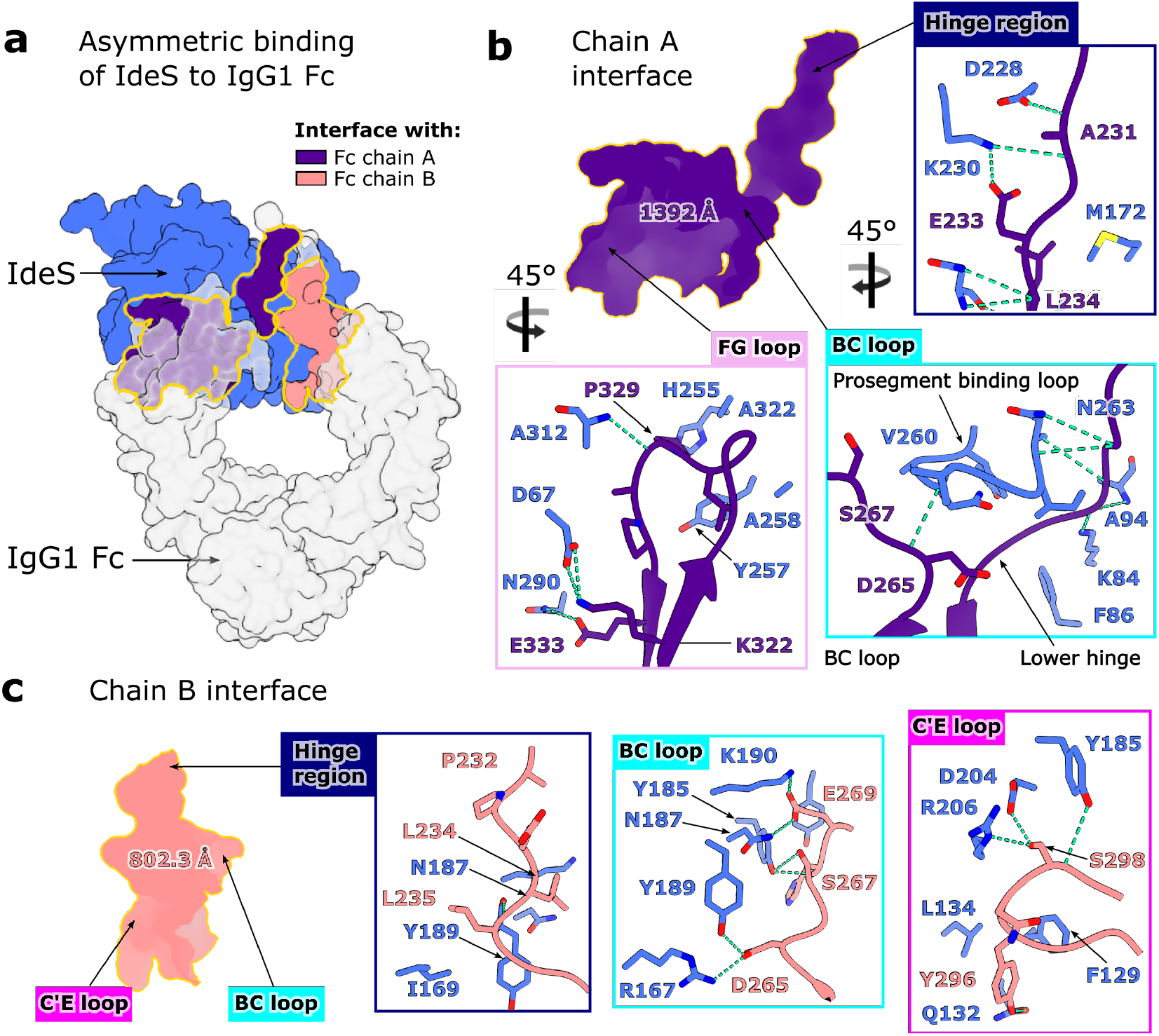
Asymmetric binding interface of IdeS-IgG1 Fc complex. **a** Overall view of complex depicted as a surface, with IdeS coloured blue and IgG1 Fc coloured silver. Interfaces of IdeS with chains A and B of the Fc are coloured indigo and coral, respectively. **b** Interface between IdeS and Fc chain A, involving the Fc hinge region, BC loop and FG loop. **c** Interface between IdeS and Fc chain B, involving the hinge region, BC loop and C’E loop. Residues involved in binding are depicted as sticks and coloured by heteroatom, with hydrogen bonds depicted as green dashes.

Alanine substitution mutations within this loop were previously found to have little effect on IdeS binding to IgG, or its catalytic activity^44^. Our structure shows, however, that the majority of interactions present here involve the IdeS backbone, whose conformation won’t be significantly altered by alanine mutations. The inability of IdeS to cleave IgG hinge-mimicking peptides^30^ also indicates an occlusion of the active site in the absence of substrate, especially given the strong potential of hydrogen bonding and hydrophobic interactions observed at the Fc hinge (discussed in the following section). We therefore conclude that this loop is important for IdeS function, specifically in mediating substrate access to the active site.

#### Interaction of Fc hinge at active site

We observe clear density for the Fc hinge region bound within the active site cavity (Fig. 2c): the carbonyl oxygen of G236 is hydrogen bonded to the amide nitrogen of the catalytic cysteine (mutated to alanine here) and the side chain of K84, which collectively form the oxyanion hole, as predicted^40,44^. Binding of the hinge distorts the peptide backbone at G236 in order to promote scissile bond cleavage (Fig. 2c); this residue is thus identified in Molprobity^45,46^ as a Ramachandran outlier. Superposition of wild-type IdeS (PDB code 2AU1) gives an indication for placement of the catalytic cysteine side chain (Supplementary Fig. 3b): in this conformation, the cysteine sulphur is ideally poised for nucleophilic attack on the carbonyl carbon within the scissile peptide bond. A water molecule observed within the active site (Fig. 2c), held in position *via* hydrogen bonds to L92, G95 and V171 backbone atoms (within IdeS) and the carbonyl oxygen of L235 in the Fc hinge, is well-placed to act as a base catalyst of the emerging covalent tetrahedral intermediate.

#### Extended exosite binding to the Fc Cγ2 domains

It has long been suspected that IdeS must recognise its sole substrate IgG with exosite binding^30,40,44^. Our structure now reveals that IdeS binds across both chains of the Fc (Fig. 3a). Unsurprisingly, the most extensive interface is formed with the Fc chain being cleaved (chain A in our structure) (Fig. 3b), with an interface area of 1392 Å^2^ and a solvation free energy gain upon interface formation of -15.9 kcal/mol, as calculated by PISA^34^. The interface extends across the entire hinge region (P228-S239; Fig. 3b), with hydrogen bonds formed with A231, L234, G236 and G237 backbone atoms and the E233 side chain, and favourable hydrophobic interactions predicted here (inferred by positive solvation energies of hinge residues). Within the Fc Cγ2 domain, IdeS interacts with residues in proximity of the Fc BC loop, which aids in stabilising an “open” conformation of the prosegment binding loop (as discussed above), and additionally the FG loop (Fig. 3b).

A secondary interface is formed across chain B of the Fc (Fig. 3c), with an interface area of 802.3 Å^2^ and a solvation free energy gain of -7.7 kcal/mol. A smaller proportion of the Fc hinge contributes (A231-G237), but PISA predicts favourable hydrophobic interactions here, albeit not to the same extent as chain A. Subsequent recognition of this Fc chain is driven by interactions with the BC loop, and, in contrast to chain A, the C’E loop containing the N-linked glycan (Fig. 3c). PISA additionally predicts a small number of interactions between the enzyme and the Fc N-linked glycans; the lack of electron density for any monosaccharides past β-1,2-linked GlcNAc suggests that any further glycan processing doesn’t affect complex formation, and that IdeS can accommodate IgG with heterogenous glycosylation.

Although IdeS interacts with both chains in the Fc hinge simultaneously, following cleavage of the first chain, the complex would need to dissociate before the second cleavage could occur. This observation is also evidenced by detection of single-cleaved Fc in enzymatic assays and in clinical studies^8,36,47,48^. We suspect that the binding interface is altered for single-cleaved Fc and that this explains its slower rate of cleavage^8,9,36^. It is also interesting to note that, aside from the hinge region, IdeS binds Fc regions also implicit in FcγR binding, an observation also inferred by its ability to counteract Fc-mediated effector functions by competitive binding inhibition^6^. Moreover, we observe that IdeS residues interacting with the Fc are largely conserved across the two IdeS isoforms, and any substitutions are mostly to similar amino acids, which aids in explaining their near identical activity^7^.

### The EndoS-IgG1 Fc complex

To date, there are several known structures of endoglycosidases in complex with their glycan substrates^26,33,49-51^. Here, we present the structure of truncated EndoS (residues 98-995, as described previously^31^) in complex with its IgG1 Fc substrate (Fc E382R mutant). A catalytically inactive version of EndoS was generated by the inclusion of D233A/E235L substitutions, as described previously^33^. The complex crystallised in space group *P*2_1_2_1_2_1_ and was refined to a resolution of 3.2 Å (Supplementary Table 3); the final structure is depicted in Fig. 4.

**Fig. 4:**
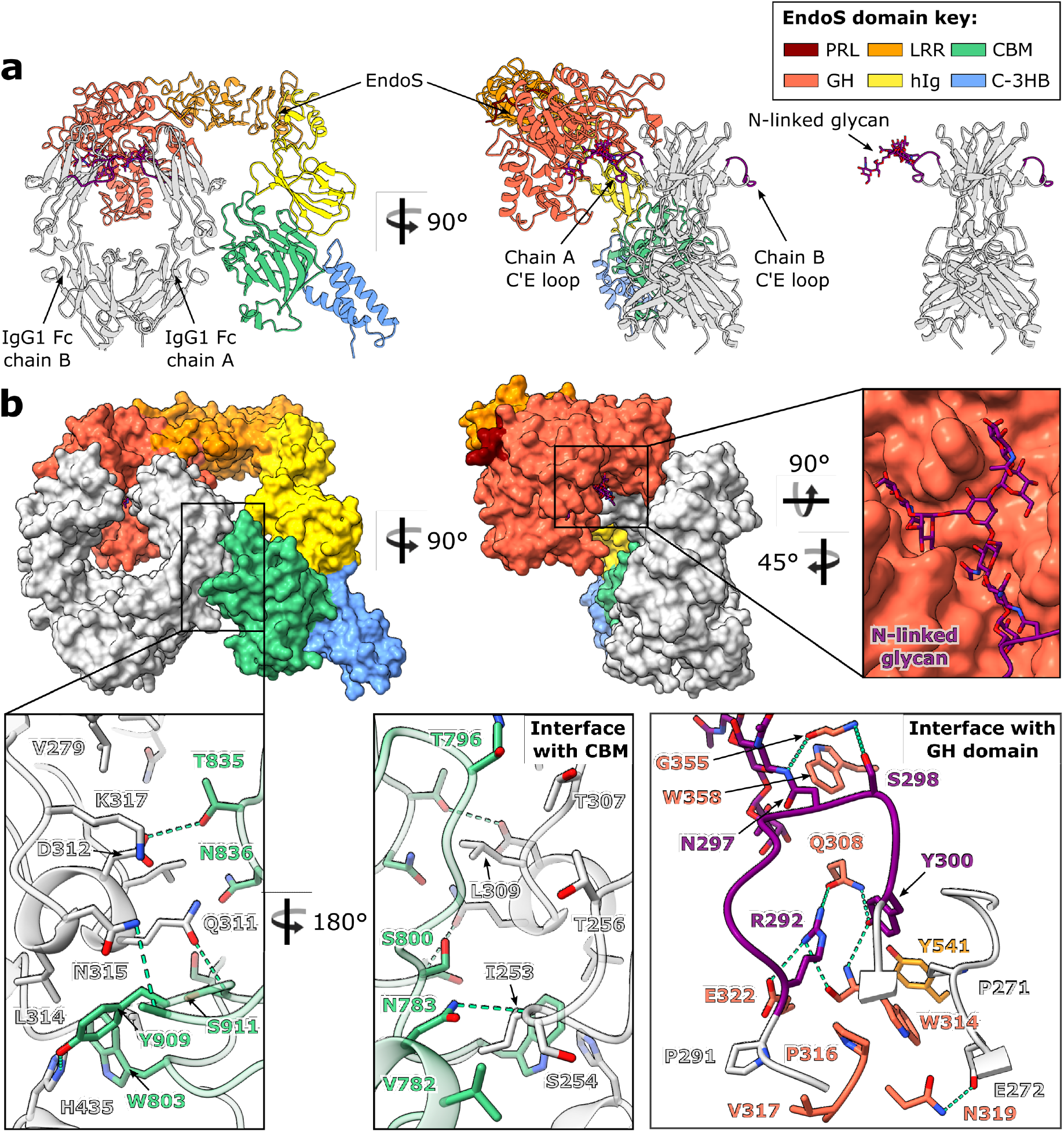
Crystal structure of EndoS^D233A/E235L^-IgG1 Fc^E382R^ complex. **a** Overall structure of complex depicted as a cartoon. IgG1 Fc is coloured silver, with its C’E loops coloured purple; the N-linked glycan is shown as sticks and coloured by heteroatom. EndoS domains are coloured as follows: proline-rich loop (PRL), maroon; glycosidase domain (GH), red; leucine-rich repeat domain (LRR), orange; hybrid Ig domain (hIg), yellow; carbohydrate-binding module (CBM), green; C-terminal 3-helix bundle (C-3HB), blue. **b** EndoS-Fc complex depicted as a surface, highlighting binding to IgG1 Fc by the CBM and GH domains. Residues involved in binding are depicted as sticks and coloured by heteroatom. Hydrogen bonds are depicted as green dashes.

#### Overall structure of EndoS-IgG1 Fc complex

Our structure of EndoS shows the same “V” shape as observed in its previously solved structures^31,33^, comprising, from the N- to the C terminus: a proline-rich loop (residues 98-112) a glycosidase domain (residues 113-445), a leucine-rich repeat domain (residues 446-631), a hybrid Ig domain (residues 632-764), a carbohydrate-binding module (CBM; residues 765-923) and a C-terminal three-helix bundle domain (C-3HB; residues 924-995) (Fig. 4a). One Cγ2 domain in IgG1 Fc (chain A in our structure) binds across the termini of the “V”, in-between the glycosidase domain and CBM, with the rest of the antibody remaining exposed to the surrounding solvent. The N-linked glycan on this chain is “flipped-out” from its usually-observed position between the two Fc Cγ2 domains^52^ and is bound within the previously-identified glycosidase domain cavity^33^ (Fig. 4b). A structural overlay with full-length EndoS in complex with its G2 oligosaccharide substrate (PDB code 6EN3) shows that the overall morphology and domain organisation of EndoS is approximately maintained (Supplementary Fig. 4a), apart from a slight shift of the CBM and C-3HB, likely due to a pinching of the CBM around the Fc as it binds.

#### Role of CBM in governing specificity for IgG

Our structure of the EndoS-Fc complex reveals how one Cγ2 domain of the Fc binds across the glycosidase domain and CBM (Fig. 4). As calculated by PISA^34^, the interface between chain A of the Fc and EndoS comprises an area of 1356.8 Å^2^ and yields a solvation free energy gain of -9.5 kcal/mol. The glycosidase domain of EndoS is observed forming contacts with the glycan-containing C’E loop, while the CBM forms additional interactions at the Fc Cγ2-Cγ3 interface (Fig. 4b). We note that residue W803 within the CBM, whose substitution to an alanine has previously been shown to abolish hydrolytic activity against all human IgG subclasses^31^, appears to act as a hydrophobic “plug”: it binds within a cavity at the Cγ2-Cγ3 interface containing Fc residues I253, H310, L314 and H435 (Fig. 4b), and has the highest solvation energy (of 2.02 kcal/M) of all EndoS residues calculated by PISA, indicating that strong hydrophobic interactions are present here. A small number of contacts is also predicted between EndoS and the second Cγ2 domain, although these are unlikely to be necessary for complex formation, given that EndoS can cleave the Fc Cγ2 lacking the hinge region (likely monomeric)^32^.

The complex structure presented here corroborates previous findings that both the glycosidase domain and the CBM are important for IgG Fc binding^31^ and glycan hydrolysis^32^, and that EndoS can cleave the Cγ2 homodimer fragment of IgG Fc^32^. The related enzyme EndoS2^25^ likely binds IgG in a similar manner: hydrogen-deuterium exchange mass spectrometry on this complex has similarly indicated strong binding of IgG to the glycosidase domain and the CBM^26^. While mutation of residues within the glycan binding site of both enzymes completely abolishes their hydrolytic activity^26,33^, EndoS lacking the CBM can still hydrolyse IgG, albeit at greatly reduced capacity^31,32^. Therefore, the CBM appears to drive additional specificity of EndoS for the Fc peptide surface.

Interestingly, although previous work has indicated that it can bind galactose (albeit with low affinity)^32^, the CBM doesn’t bind carbohydrate, and the N- and C-terminal 3 helix bundles, which are homologous to IgG-binding protein A from *Staphylococcus aureus*^33,53^, don’t bind protein. A structural overlay of complexed EndoS with full-length EndoS (PDB code 6EN3) indicates that the N-terminal bundle would not contact the Fc (Supplementary Fig. 4a), thus its contribution to EndoS-IgG binding and glycan hydrolysis is likely solely due to stabilisation of the glycosidase domain, as suggested previously^33^. Indeed, existence of the crystal structure is evidence in itself that EndoS forms a stable complex with IgG in its absence.

#### Stoichiometry of EndoS-Fc complex

Within the crystal, we observe a 2:1 stoichiometry of EndoS binding to IgG Fc in the complex (Fig. 5a). The N297 glycan within chain A of the Fc binds one EndoS molecule, while its counterpart in chain B, although not fully visible in the electron density, appears to be bound to a second EndoS molecule present in the asymmetric unit of the crystal (Fig. 5a). We observe clear electron density for the chain A glycan binding within the EndoS glycosidase domain cavity previously identified^33^, and for the C’E loop which covalently links the glycan to the Fc peptide (Fig. 5). This observation of an Fc glycan in this conformation is in strong contrast to typical crystal structures of IgG Fc, whose N-linked glycans are found between the Cγ2 domains^52^ (Fig. 5b).

**Fig. 5:**
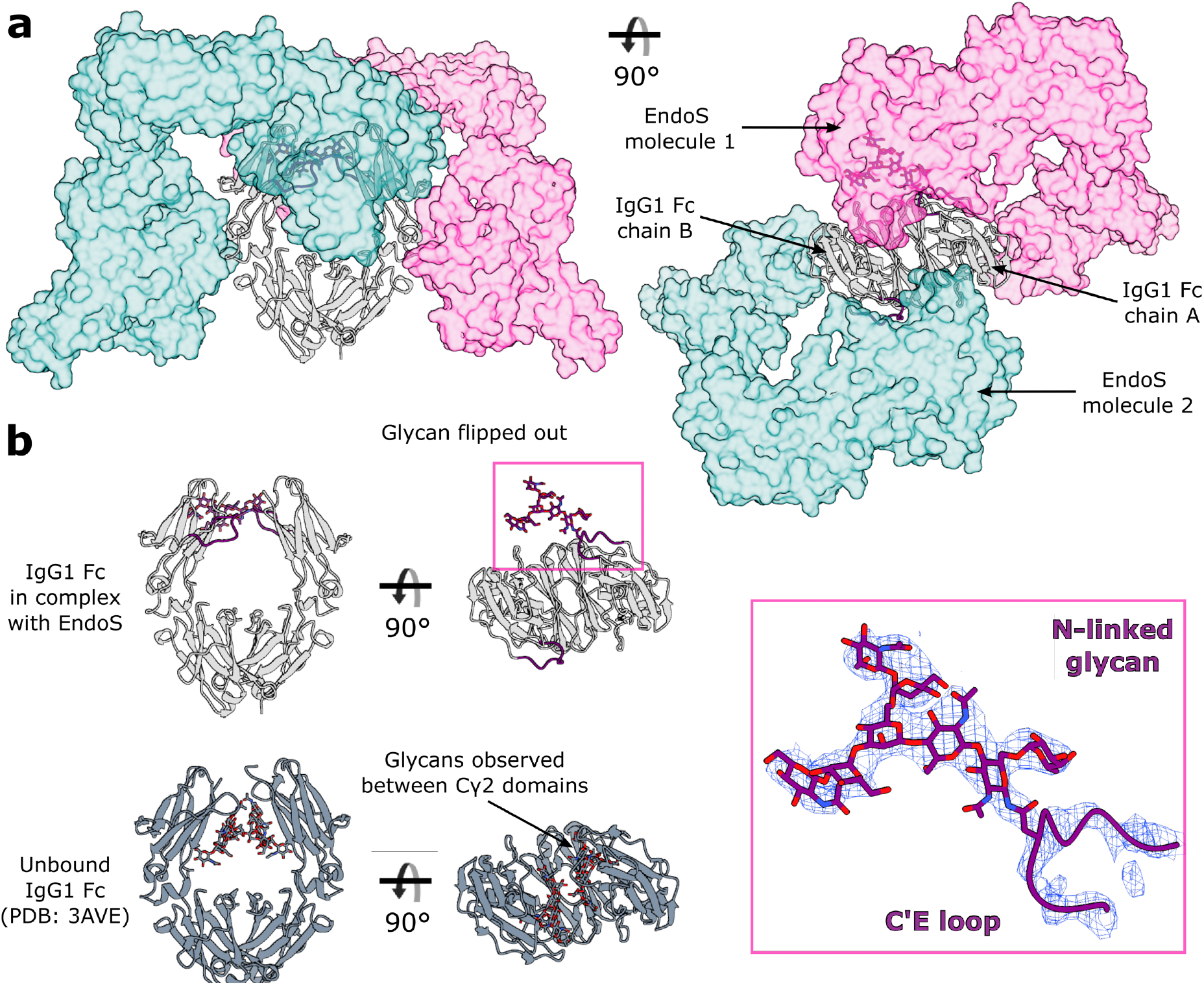
Stoichiometry of EndoS-IgG Fc complex. **a** Two copies of EndoS bound to a single IgG1 Fc, as observed in the crystal structure. The two EndoS molecules are coloured teal and pink and depicted as a surface at 50 % transparency, while the Fc is coloured silver and depicted as a cartoon. C’E loops within the Fc are coloured purple; the N-linked glycan is depicted as sticks and coloured by heteroatom. **b** Comparison of N-linked glycan positions observed in IgG1 Fc bound to EndoS, and a wild-type IgG1 Fc structure (PDB code 3AVE, coloured in dark grey). N-linked glycan is observed in a “flipped-out” structure in the complexed Fc, while N-linked glycans in typical Fc structures are observed between the Fc Cγ2 domains. Electron density from the final 2F_obs_-F_calc_ map, corresponding to the glycan and C’E loop, is shown (weighted at 1.1 σ).

It is fascinating to observe the glycan trapped in this “flipped-out” conformation, and this substantiates several recent studies documenting the existence of IgG Fc glycan conformational heterogeneity^54-58^. Superposition of this complexed IgG with a wild-type Fc (PDB code 3AVE) illustrates that movement of the glycan into this position is governed by movement of the C’E loop only (Supplementary Fig. 4b), although it is possible that the lower resolution of the data is masking small chain shifts. Moreover, it appears that the capture of Fc N-linked glycans in this state allows space for two enzymes to bind simultaneously; however, there is no evidence to suggest that this 2:1 assembly is required for activity, especially given previous work showing that EndoS is largely monomeric in solution^31,33^. Although EndoS crystallised here is lacking the N-terminal 3-helix bundle, a structural superposition with full-length EndoS (Supplementary Fig. 4a) suggests 2:1 binding would be able to occur in its presence.

### Perspectives

The crystal structures presented here provide a structural rationale for the unique properties of these two enzymes, particularly their exquisite substrate specificity towards human IgG. Understanding the molecular basis of this activity is critical for expanding their clinical and biotechnological use. For example, the deactivation of serum IgG using both IdeS and EndoS can strengthen the potency of therapeutic antibodies^21,22^; this strategy could be applied to potentiate any therapeutic antibody, in theory, if the antibody were designed to be resistant to cleavage by these enzymes, a venture which can be aided greatly with structural information. This will also be invaluable in the synthesis of immunologically-distinct enzyme variants which retain identical activity, for their long-term therapeutic use. While EndoS variants have already been designed to expand the ability to engineer antibody glycosylation^27-29^, the structural information presented here will allow this to be extended further. To conclude, this work will assist in the continued development of IdeS and EndoS as enzymatic tools with wide clinical and biotechnological applications.

## Materials and methods

### Cloning, Protein Expression and Purification

#### IdeS/EndoS

Gene fragments encoding IdeS^C94A^ (amino acids 41-339, gene accession number AFC66043.1) and EndoS^D233A/E235L^ (amino acids 98-995, as described previously^31,33^) were synthesised to contain a C-terminal linker and His tag (sequence LEHHHHHH), and cloned into pET21a(+) vectors by NBS Biologicals. Constructs were expressed in *E. coli* BL21 (DE3) cells. Cells were grown at 37 °C in Terrific Broth (Melford) in the presence of 100 mg/mL ampicillin and 34 mg/mL chloramphenicol, until an OD_600_ of 0.8 was reached, when protein expression was induced by addition of 1 mM IPTG. Cells were left to shake overnight at 25 °C, 200 rpm (Innova 43R incubator, New Brunswick Scientific). Cells were collected by centrifugation at 6,220 x *g* for 20 minutes, resuspended in PBS containing 2 μg/mL DNAse1 (Sigma) and a pinch of lysozyme (Sigma), homogenised using a glass homogeniser and broken apart using a cell disruptor (Constant Cell Disruption Systems). The remaining sample was centrifuged first at 3,100 x *g* for 20 minutes, then again at 100,000 x *g* for one hour, to remove remaining cell debris and cell membranes. The resulting supernatant was subsequently filtered through a 0.2 μm membrane. Proteins were purified from the supernatant using Ni affinity chromatography with a HisTrap HP column (Cytiva) followed by size exclusion chromatography with a Superdex 75 16/600 column (Cytiva), equilibrated in 10 mM HEPES, 150 mM NaCl, pH 8.0.

#### IgG1 Fcs

E382R/S/A mutations were introduced into IgG1 Fc encoded within a pFUSE-hIgG1-Fc vector using site-directed mutagenesis (QuikChange II kit, Agilent), using mutagenic primers (Supplementary Table 4) synthesised by Eurofins Genomics. Fcs were transiently expressed in FreeStyle293F cells (ThermoFisher) by incubating at 37 °C, 8 % CO_2_ and shaking at 125 rpm (New Brunswick S41i incubator), and harvested after seven days by centrifugation at 3,100 x *g* for 30 minutes. Supernatant was filtered through a 0.2 μm membrane and antibodies purified by affinity purification with a HiTrap Protein A HP column (Cytiva), followed by size exclusion chromatography with a Superdex 200 16/600 column (Cytiva) in 10 mM HEPES, 150 mM NaCl (pH 8.0).

### Protein Crystallisation

Protein concentrations were determined with a DS-11+ Spectrophotometer (DeNovix), using molecular weight and extinction coefficients calculated by the ProtParam tool^59^. IdeS/EndoS were combined with IgG1 Fcs in a 1:1 molar ratio and applied to a Superdex 200 16/600 column (Cytiva) equilibrated in 10 mM HEPES, 150 mM NaCl (pH 8.0). Fractions corresponding to the main peak only were pooled for crystallisation. Purified complexes were exchanged into 50 mM HEPES, 150 mM KCl (pH 7.5) prior to crystallization, using a Vivaspin 20 centrifugal concentrator. Sitting drop vapour diffusion crystallisation trays were set up using an Oryx4 robot (Douglas Instruments). Crystals of the IdeS-Fc complex were grown in 0.12 M monosaccharides mix, 0.1 M buffer system 3 (pH 8.5), 30 % v/v precipitant mix 1 (Morpheus crystallization screen, Molecular Dimensions). Crystals of the EndoS-Fc complex were grown in 0.09 M halogens, 0.1 M buffer system 2 (pH 7.5), 37.5 % v/v precipitant mix 4 (Morpheus crystallization screen, Molecular Dimensions). Crystals of IgG1 Fc E382S was grown in 0.2 M ammonium sulphate, 0.1 M Tris (pH 7.5), 25 % w/v PEG 8000. Crystals were cryo-protected in mother liquor with 20 % glycerol added and flash-frozen in liquid nitrogen.

### Data collection and Structure Determination

Data collection was carried out at the European Radiation Synchrotron Facility (Grenoble, France) for IdeS-Fc and Fc E382S (beamline ID30A-3), and Diamond Light Source (Oxford, UK) for EndoS-Fc (beamline I03). Data processing of diffraction images was carried out using DIALS^60^ and XDS^61^. Structures were solved by molecular replacement with the program Molrep^62^. 3AVE was used as a search model to solve the IgG1 Fc^E382S^ structure; IdeS-Fc and EndoS-Fc were solved using initial search models for the enzyme (PDB code 1Y08 for IdeS; 6EN3 for EndoS), after which the resulting solution was used as a fixed model for a second round of molecular replacement, using 3AVE as the search model. Models were improved with successive rounds of model building and refinement, using Coot^63^ and Refmac5^64^, respectively, within the ccp4i2 suite^35^. Due to the presence of twinning in the IdeS-Fc data, this structure was refined with the option for twinning ticked in Refmac5. All structures were refined using local non-crystallographic symmetry restraints. PDB-REDO^65^ was used to generate restraints for the IgG1 Fc^E382S^ model for use in refinement. MolProbity^46^ and the PDB validation server^66,67^ were used for model validation prior to deposition. Carbohydrates were modelled in Coot^68^ and validated using Privateer^69^. Protein complex interfaces were analysed using PISA^34^. UCSF ChimeraX^37^ was used to prepare figures depicting protein structure.

## Data Availability

Structure factor files and atomic coordinates for each of the models presented in this study have been deposited in the PDB, with accession codes 8A47 (IdeS-Fc), 8A48 (IgG1 Fc^E382S^) and 8A49 (EndoS-Fc).

## Acknowledgements

We are grateful to the beamline scientists on I03 at Diamond Light Source (DLS) and ID30-A3 at the European Synchrotron Radiation Facility (ESRF), and to Chris Holes for his support with the macromolecular crystallization facility at the University of Southampton. This work was supported by DLS, ESRF (MX2373) and the School of Biological Sciences, University of Southampton.

