## Supplementary Tables 1-4, Supplementary Figures 1-4 for "Extensive substrate recognition by the streptococcal antibody-degrading enzymes IdeS and EndoS"

### SUPPLEMENTARY INFORMATION

| | |
|---|---|
| **Data Collection** | |
| Beamline | ID30A-3 (European Synchrotron Radiation Facility) |
| Resolution range (Å) | 47.5-3.04 |
| Space group | $P3_221$ |
| Unit cell dimensions:<br>*a, b, c* (Å)<br>α,β,γ (degrees) | <br>106.78, 106.78, 104.01<br>90, 90, 120 |
| Wavelength (Å) | 0.968 |
| Unique reflections | 13476 (2358) |
| Completeness (%) | 99.7 (98.4) |
| $R_{merge}$ (%) | 0.034 (1.117) |
| $R_{meas}$ (%) | 0.048 (1.580) |
| $R_{pim}$ (%) | 0.034 (1.117) |
| I/σ(I) | 15.2 (0.6) |
| Multiplicity | 1.9 (1.9) |
| **Refinement** | |
| Number of reflections (all/free) | 13476/654 |
| $R_{work}$ (%) | 19.7 |
| $R_{free}$ (%) | 22.0 |
| RMSD:<br>Bonds (Å)<br>Angles (degrees) | <br>0.0027<br>0.999 |
| Molecules per ASU | 1 |
| Atoms per ASU | 3,565 |
| Average B factors ($Å^2$) (protein/ligand/water) | (141.52/177.82/107.72) |
| Model quality (Ramachandran plot):<br>Most favoured region (%)<br>Allowed region (%) | <br>96.37<br>3.15 |

**Supplementary Table 1: Crystallographic data collection and refinement statistics for less-crystallisable "Fx" E382S mutant.** Values for the highest resolution shell are shown in parentheses.

| **Data Collection** | |
|---|---|
| Beamline | ID30A-3 (European Synchrotron Radiation Facility) |
| Resolution range (Å) | 48.53 – 2.34 (2.38-2.34) |
| Space group | *C* 1 2 1 |
| Unit cell dimensions:<br>*a, b, c* (Å)<br>α,β,γ (degrees) | <br>217.56, 108.45, 63.05<br>90.00, 90.06, 90.00 |
| Wavelength (Å) | 0.9677 |
| Unique reflections | 60351 (3005) |
| Completeness (%) | 97.7 (98.8) |
| $R_{merge}$ (%) | 22.9 (87.7) |
| $R_{meas}$ (%) | 25.5 (97.8) |
| $R_{pim}$ (%) | 11.2 (42.6) |
| I/σ(I) | 5.3 (1.0) |
| Multiplicity | 4.96 (4.98) |
| **Refinement** | |
| Number of reflections (all/free) | 60350/3049 |
| $R_{work}$ (%) | 18.2 |
| $R_{free}$ (%) | 20.6 |
| RMSD:<br>Bonds (Å)<br>Angles (degrees) | <br>0.0033<br>0.965 |
| Molecules per ASU | 2 |
| Atoms per ASU | 6,192 |
| Average B factors ($Å^2$) (protein/ligand/water) | (38.70/40.08/38.95) |
| Model quality (Ramachandran plot):<br>Most favoured region (%)<br>Allowed region (%) | <br>96.82<br>2.9 |

**Supplementary Table 2: Crystallographic data collection and refinement statistics for IdeS-IgG1 Fc complex.** Values for the highest resolution shell are shown in parentheses.

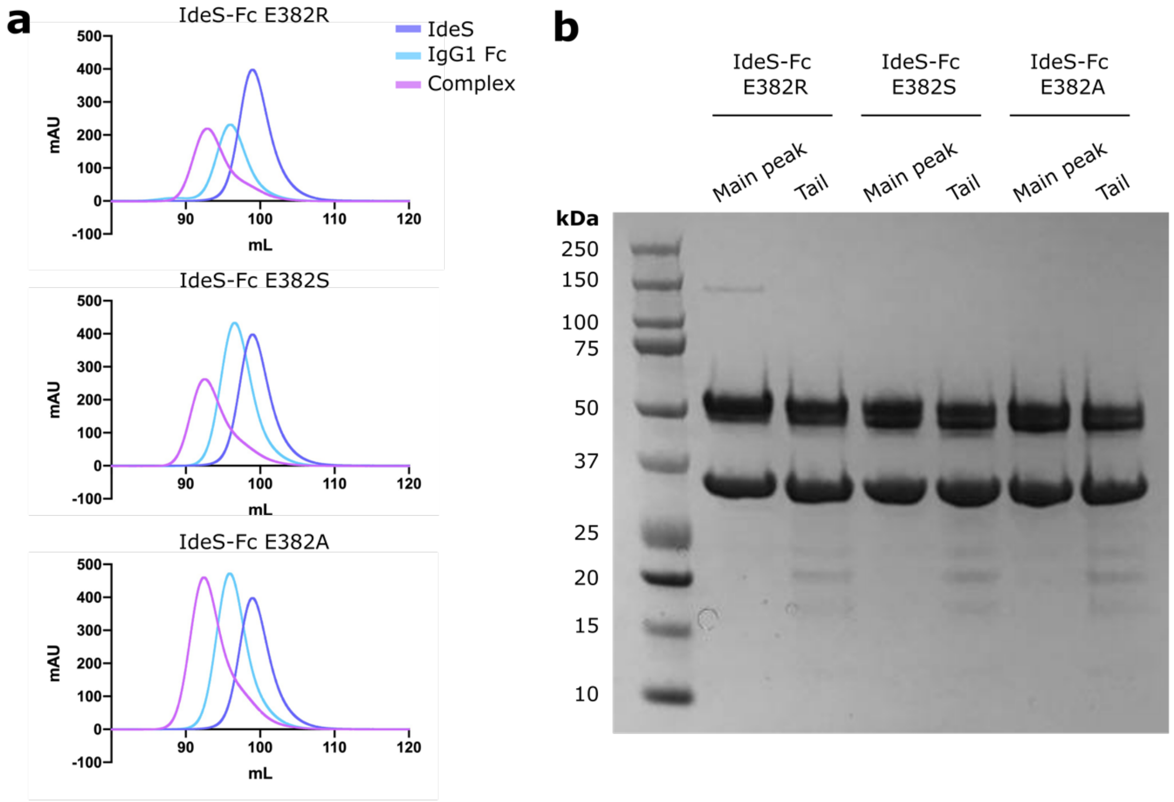


**Supplementary Fig. 1: Purification of IgG1 Fc-IdeS complexes. a** Analytical size exclusion chromatography (SEC) of IdeS-IgG1 Fc complexes. IdeS and Fcs containing E382R, E382S and E382A mutations were first applied to a Superdex 16/600 s200 column, then complexes were combined in a 1:1 molar ratio and applied to the column again. **b** SDS-PAGE of the main peak and tail fractions following purification of each complex. Main peaks for each complex contain dominant bands for Fc (~50 kDa) and IdeS (~34 kDa). Main pean fractions only were taken forward for crystallization.

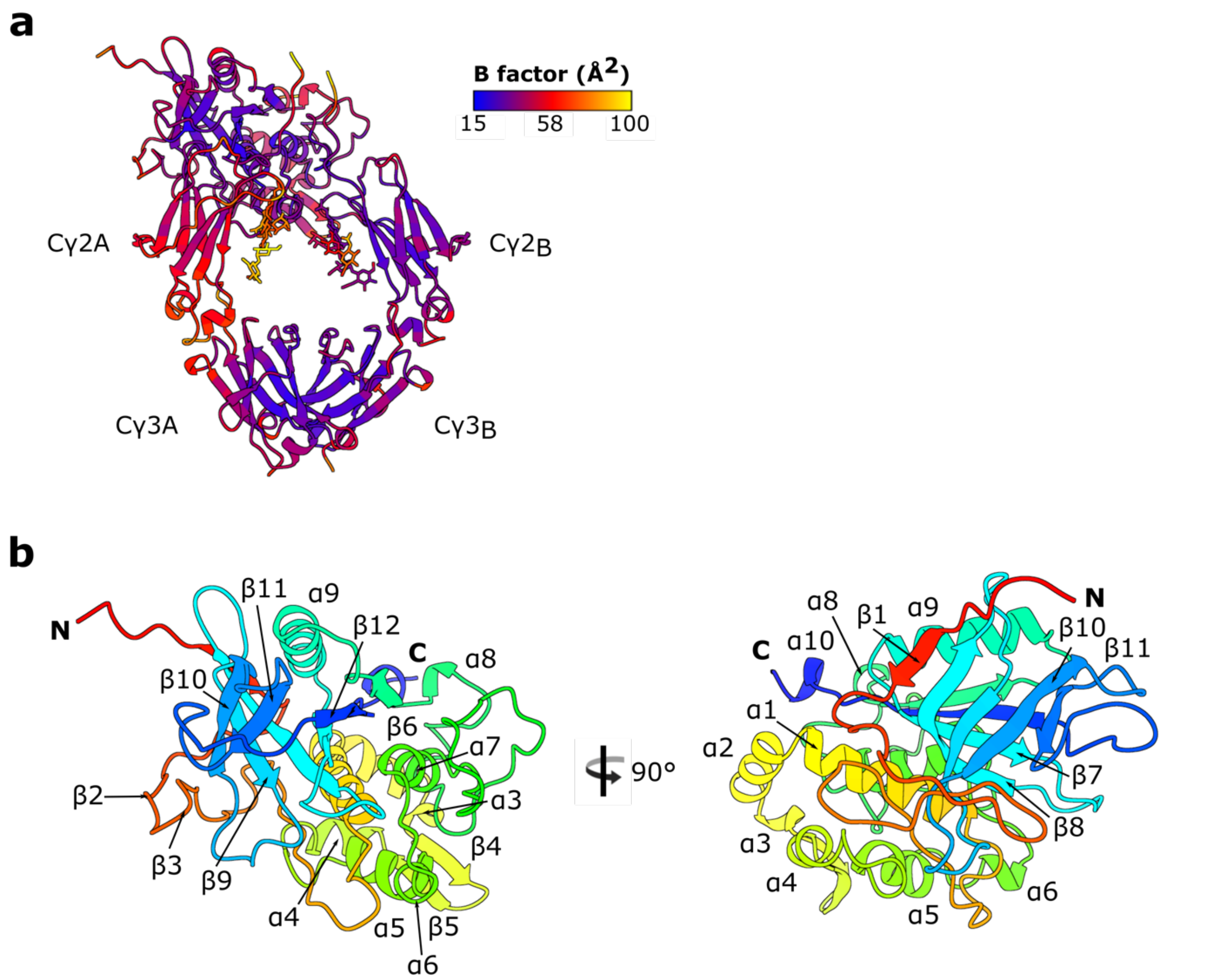


**Supplementary Fig. 2: Analysis of IdeS-Fc crystal structure. a** Distribution of average B factors per residue for IdeS-Fc complex. **b** Front and side views of complexed IdeS, coloured as a rainbow from the N- (red) to C-terminus (blue) and labelled with secondary structure, as calculated by DSSP[1,2].

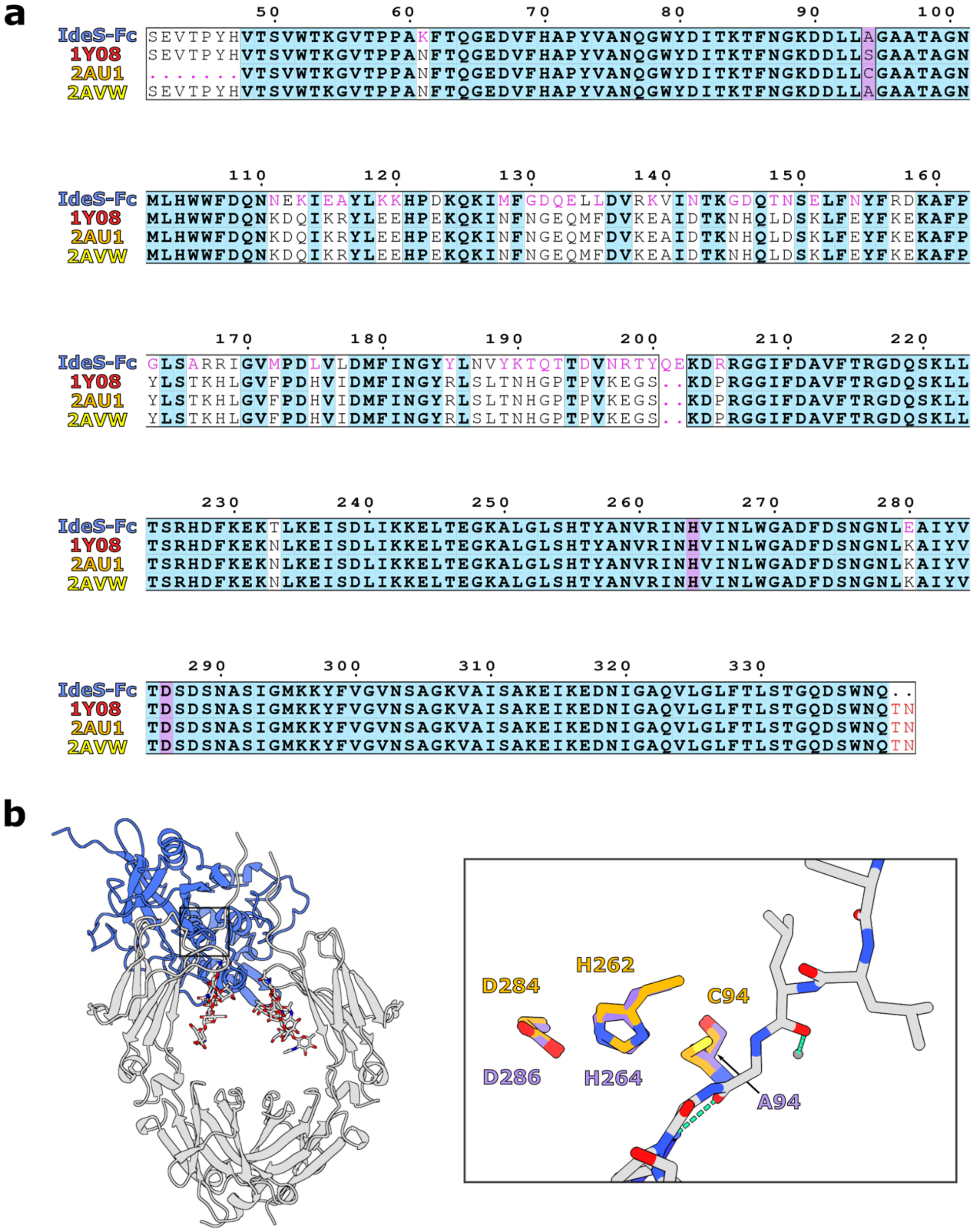


**Supplementary Fig. 3: Analysis of IdeS-Fc crystal structure (continued). a** Multiple sequence alignment of complexed IdeS vs published apo IdeS structures (PDB codes 1Y08, 2AU1 and 2AVW), generated in Clustal Omega[3] and depicted using ESPript 3.0[4]. Conserved residues are coloured blue; catalytic triad residues are highlighted in purple; similar residues are shown in black font; residues with differing chemical properties are coloured in pink font. Numbering is shown for the complexed form of IdeS (Mac-2). **b** Superposition of catalytic residues of complexed IdeS vs apo wild-type IdeS (PDB code 2AU1). IgG1 Fc is coloured silver.

| **Data Collection** | |
|---|---|
| Beamline | I03 (Diamond Light Source) |
| Resolution range (Å) | 49.78-3.2 |
| Space group | $P2_12_12_1$ |
| Unit cell dimensions:<br>*a, b, c* (Å)<br>α,β,γ (degrees) | <br>96.529, 174.294, 193.059<br>90, 90, 90 |
| Wavelength (Å) | 0.9763 |
| Unique reflections | 54582 |
| Completeness (%) | 100 |
| $R_{merge}$ (%) | 0.177 |
| $R_{meas}$ (%) | 0.184 |
| $R_{pim}$ (%) | 0.049 |
| I/σ(I) | 9 |
| Multiplicity | 13.9 |
| **Refinement** | |
| Number of reflections (all/free) | 54513/2764 |
| $R_{work}$ (%) | 25.8 |
| $R_{free}$ (%) | 30.4 |
| RMSD:<br>Bonds (Å)<br>Angles (degrees) | <br>0.0020<br>0.616 |
| Molecules per ASU | 3 |
| Atoms per ASU | 17339 |
| Average B factors ($Å^2$) (protein/ligand/water) | 157.07, 163.91, 89.09 |
| Model quality (Ramachandran plot):<br>Most favoured region (%)<br>Allowed region (%) | <br>93.27<br>5.85 |

**Supplementary Table 3: Crystallographic data collection and refinement statistics for EndoS-Fc complex.** Values for the highest resolution shell are shown in parentheses.

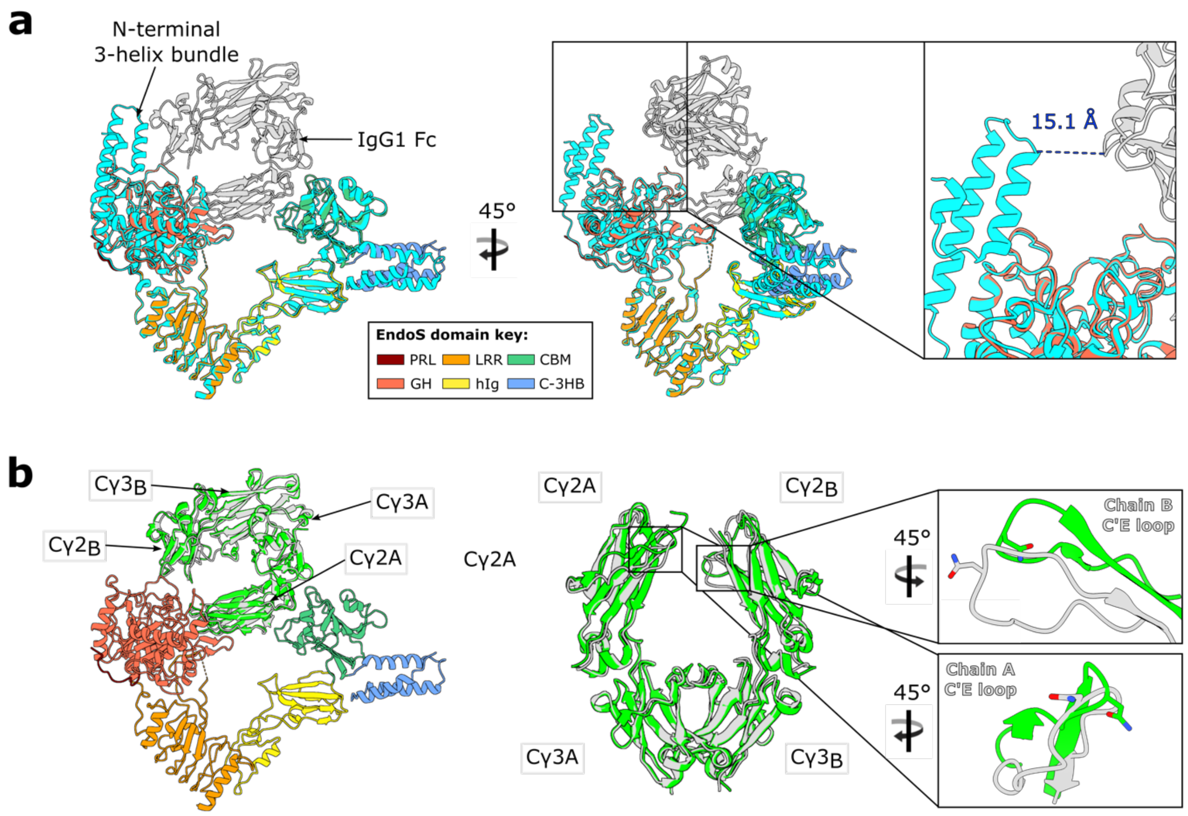


**Supplementary Fig. 4: Superposition of EndoS-Fc complex with respective apo structures of each component. a** Superposition of full-length EndoS (PDB code 6EN3, coloured in cyan) with complexed EndoS. Focused view of N-terminal 3-helix bundle (not present in the complexed EndoS construct) indicates that it would not bind IgG1 Fc. **b** Superposition of wild-type IgG1 Fc (PDB code 3AVE; coloured in lime) with complexed IgG1 Fc, which reveals conformational changes in the C′E loops containing the N-linked glycans at N297 (whose side chain is shown as sticks and coloured by heteroatom). **a,b** Complexed EndoS and Fc are coloured as in Fig. 4a.

| IgG1 Fc construct | Mutagenic primer sequence (5′ → 3′) | |
|---|---|---|
| | **Forward** | **Reverse** |
| E382R | GACATCGCCGTGGAGTGGAGTGGAG GAGCAATGGGCAGCCGGAGAACAAC | TTGTTCTCCGGCTGCCCATTGCTCCTCC ACTCCACGGCGATGTCG |
| E382S | GACATCGCCGTGGAGTGGAGTGGAG CAGCAATGGGCAGCCGGAGAACAAC | TTGTTCTCCGGCTGCCCATTGCTGCTCC ACTCCACGGCGATGTCG |
| E382A | GACATCGCCGTGGAGTGGAGTGGGC GAGCAATGGGCAGCCGGAGAACAAC | TTGTTCTCCGGCTGCCCATTGCTCGCC CACTCCACGGCGATGTCG |

**Supplementary Table 4: Primers used for site-directed mutagenesis of IgG1 Fc constructs.** Primers are written 5′ to 3′.
